# The effects of DNA repair factor-Cas9 fusions on homing gene drive performance

**DOI:** 10.64898/2026.10.20.760381

**Authors:** Yingke Wu, Xuejiao Xu, Nan Liang, Yunchen Xia, Andrea Y.N. Tan, Jie Yang, Xiaozhen Yang, Jackson Champer

**Author notes:** Equal contribution (Yingke Wu, Xuejiao Xu).

## Abstract

CRISPR-based homing gene drives achieve super-Mendelian inheritance by converting wild-type alleles into drive alleles through homology-directed repair (HDR). However, competing end-joining repair can generate resistance alleles that limit drive spread. Although fusing Cas9 to DNA repair-associated proteins has improved HDR in conventional genome editing systems, its effectiveness in gene drives remains unclear. Here, we systematically evaluated in *Drosophila melanogaster* eight Cas9 fusion constructs previously reported to improve HDR in other systems. Although none of the fusion constructs significantly increased HDR-associated signals in S2 cells, several candidates retained relatively high activity comparable to standard Cas9. These candidates were then tested *in vivo* across multiple split drive systems, regulatory elements, target loci, and parental sexes. Cas9-POLD3 significantly increased female drive inheritance in contexts with low baseline efficiency or substantial nonfunctional resistance formation, likely by increasing effective cutting rate and potentially also by biasing repair outcomes toward drive conversion. Cas9-Geminin degron generally maintained germline drive performance while reducing embryo resistance. Together, these findings identify Cas9-POLD3 and Cas9-Geminin degron as complementary, context-specific tools for enhancing drive conversion and limiting embryo resistance, respectively. These results highlight the importance of matching fusion modules to the regulatory and genetic context of individual drive systems.

## Introduction

Gene drives offer a powerful tool to manipulate populations by preferentially spreading specific traits. They can be applied to suppress or modify pest populations^1–4^. Although many types of gene drive systems have been developed, CRISPR-based homing drives^5^ are among the most powerful approaches. In these systems, the drive allele encodes gRNAs that direct Cas9 to cleave the homologous wild-type allele. The resulting double-strand break can then be repaired by homology-directed repair (HDR) using the drive-containing chromosome as a template. This process converts drive/wild-type heterozygotes into drive homozygotes and promotes drive inheritance in offspring above the Mendelian expectation. A higher HDR-derived drive conversion rate can therefore accelerate drive spread in a population. For suppression drives, it also allows higher suppressive power to be achieved.

HDR is a template-dependent pathway that enables precise repair of Cas9-induced cleavage when a homologous donor template or sister chromatid is available, which is restricted to the S/G2 phases of the cell cycle. After cleavage, DNA ends are resected to generate 3′ single-stranded DNA, followed by homology search, strand invasion, and repair synthesis. These steps allow accurate sequence copying, but HDR is usually limited by donor or homologous template availability and requires more processing time^6^. By contrast, the competing end-joining pathway does not require a long homologous template and is active across broader cellular contexts. In this pathway, DNA ends are protected from resection and subsequently processed for ligation, providing a faster repair route^6^. However, end-joining often introduces insertions or deletions at the target site, which can prevent Cas9/gRNA recognition, thus generating resistance alleles in CRISPR-based homing drive systems^6,7^.

Low drive conversion rate and resistance allele formation are therefore major challenges that limit the spread of homing drive systems^8–11^. Efficient drive conversion depends on sufficient Cas9 expression in the germline during the repair window favorable for HDR^5,12^. Resistance alleles can be generated in the germline when cleavage is repaired by end-joining, or in embryos through cleavage caused by maternally deposited Cas9, where desired drive conversion does not occur even if a template is available. These resistance alleles prevent successful drive copying and weaken drive performance. Several strategies have been explored to increase HDR efficiency and reduce resistance allele formation, including Cas9 promoter optimization^10,13–15^ and gRNA multiplexing^16^. However, for some non-model and non-*Anopheles* species, optimal Cas9 promoters are difficult to identify because their germline biology is less well understood and may vary among species^13,14,17^. Although gRNA multiplexing may moderately increase homing rate (compared to one gRNA in some circumstances) and substantially reduce functional resistance allele formation, it does not by itself eliminate nonfunctional resistance alleles, whose consequences depend on the target gene and drive design and are particularly detrimental to suppression drives ^16,18,19^. These limitations suggest that additional strategies are needed to improve HDR and reduce end-joining mediated resistance formation.

Several strategies have been developed to improve HDR efficiency in conventional CRISPR gene editing systems. One approach is to recruit the DNA donor to the Cas9/gRNA complex, thereby increasing the local availability of the donor template for repair. For example, ssODNs can be covalently tethered to the Cas9-guide complex through a fused HUH endonuclease^20^. However, these approaches are difficult to adapt to an *in vivo* gene drive context, where the double-strand DNA template is provided by the homologous chromosome. Another option is to disrupt proteins involved in end-joining repair, such as Ku70/Ku80. Although this strategy can enhance HDR in some species, disrupting these genes *in vivo* may cause substantial fitness costs^6,7^ and limit its feasibility for gene drives. A previous study also showed little benefit when these genes were disrupted by RNAi^21^.

One attractive strategy is to tether DNA repair-associated proteins directly to Cas9. Several Cas9 fusion proteins have been reported to increase the frequency or precision of knock-in and transgene integration. These Cas9 fusion designs can be broadly grouped into four classes. The first class includes factors involved in DNA end resection, such as CtIP^22^ and MRE11^23^. These proteins promote the initiation of resection, an important step that suppresses end-joining repair and creates conditions favorable for HDR. The second class includes core HDR and repair-synthesis factors, such as POLD3^24^ and RAD51^25^. POLD3 can accelerate repair and improve HDR in a locus- and cell-type-dependent manner^24^, whereas RAD51 promotes presynaptic filament formation, homologous strand search, and strand invasion^25^. The third class includes proteins associated with epigenetic modification, such as PRDM9 and SETD2^26^. These proteins can locally deposit histone marks, including H3K4me3 and H3K36me3, which are associated with a chromatin environment more favorable for HDR. The fourth class includes cell-cycle regulatory modules, such as the Geminin degron^22^, which promote Cas9 degradation during the G1 phase, thereby restricting Cas9 activity in the HDR-permissive S/G2 phases. These approaches have been shown to improve HDR efficiency in mammalian systems to varying extents, but their effectiveness has not been demonstrated in insects.

Whether DNA repair-associated Cas9 fusions can improve homing drive performance remains unresolved. A useful fusion Cas9 for gene drive engineering must meet stricter requirements than a standard knock-in efficiency enhancer. It should preserve efficient germline cleavage, bias repair toward drive conversion, and reduce germline and/or embryo resistance allele formation. Here, we systematically evaluate eight repair-associated Cas9 fusion constructs in *Drosophila melanogaster* using an S2 cell HDR assay followed by *in vivo* testing across multiple split drive systems, regulatory elements, target loci, and parental sexes. Our results show that DNA repair-associated Cas9 fusions can improve drive performance, although their effects are strongly context dependent. Cas9-POLD3 enhances drive inheritance mainly in systems with low baseline efficiency or substantial resistance formation, whereas Cas9-Geminin degron reduces embryo resistance while preserving germline performance. By contrast, other fusions generally reduced drive performance. These findings suggest that POLD3 and the Geminin degron could address different limitations of gene drive systems and may help guide the development of more stable and efficient genetic strategies for pest control.

## Methods

### Plasmid construction

Plasmids SNc9Act5G, SNc9sp2nG, SNc9sp3nG, SNc9Cyp6nG, SNc92nG, TTChsp70c9, and BHDabg1 have been reported previously^27–29^ and were modified to generate new plasmids or applied in cell transfection or transgenic line construction in this study. SNc9Act5G contains Cas9 driven by the *Act5C* promoter and with the *Act5C* 3′ UTR, together with an EGFP marker driven by the synthetic 3×P3 promoter. SNc9sp2nG, SNc9sp3nG, and SNc9Cyp6nG share overall the same configuration as SNc9Act5G, except that Cas9 expression is driven by promoters derived from *CG7213*, *CG11929*, and *Cyp6a19* respectively, with 3′ UTR/terminators derived from *nanos*. Cas9-expressing helper TTChsp70c9 contains Cas9 driven by the *Hsp70Bb* promoter. BHDcN1 contains Cas9 under the control of *nanos* regulatory elements, a 3×P3-DsRed marker, and a gRNA gene targeting *cinnabar*. BHDabg1 expresses a gRNA to target an intergenic region between *gapdh1* and *CG12824* on chromosome 2R.

Various DNA repair-associated proteins were fused to Cas9 based on the above plasmids, either positioned at its N- or C-terminus with one or two nuclei localization signals (NLSs). The fusion proteins included the human proteins CtIP, Geminin degron, POLD3, PRDM9, and Brex27; the viral protein UL12 from herpes simplex virus 1 (HSV-1); and the *Drosophila* proteins DmCtIP-RA and Spn-A. Genomic or plasmid DNA of interest was amplified using NEBNext Ultra II Q5 Master Mix. Template plasmids were digested with restriction enzymes (NEB), followed by Gibson assembly using NEBuilder HiFi DNA Assembly Master Mix (NEB). The assembled plasmids were transformed into chemically competent *E. coli* strains DH5α (TIANGEN) or XL10 (Weidi), and positive clones were confirmed via Sanger sequencing. Plasmids were subsequently extracted using the ZymoPure Midiprep Kit (Zymo Research) and verified by full-plasmid sequencing before proceeding to cell transfection or embryo injection. Plasmid maps of all new sequences are available on GitHub (https://github.com/jchamper/Fusion-Cas9).

### Cell culture and transfection

The *Drosophila* S2 cell line was obtained from the laboratory of Jian Zhu. For routine maintenance, *Drosophila* S2 cells were cultured in Schneider’s *Drosophila* Medium (BasalMedia) supplemented with 10% (v/v) fetal bovine serum (FBS; Gibco) and 1× penicillin–streptomycin–amphotericin B solution (Solarbio). Cells were maintained at 28 °C. To establish fully adherent growth, all suspended cells were removed during passaging, allowing enrichment of an adherent S2 cell population. 24 hours prior to transfection, cells were seeded in 6-well plates at ∼12.5% confluence. Transfections were carried out using TransIT-X2 (Mirus) at one-quarter of the manufacturer’s recommended reagent amount. Cas9 and gRNA plasmids were mixed at a 4:1 molar ratio for co-transfection. At 24 hours post-transfection, cells were imaged, and the medium was replaced with fresh medium containing 4 µg/mL puromycin for selection. Cells were maintained under selection until all non-transfected cells were eliminated and the transfected population stabilized. The puromycin concentration was then reduced to 2 µg/mL as a maintenance dose to preserve population stability for subsequent experiments.

### Genomic DNA extraction and quantitative PCR

Cells were harvested by TrypLE Express (Gibco) treatment and centrifuged at 500×g for 1 minute. After removal of the supernatant, nucleic acids were extracted from the cell pellet using the AllPure DNA/RNA/Protein Kit (Magen) according to the manufacturer’s instructions.

Quantitative PCR was performed using Taq Pro Universal SYBR qPCR Master Mix (Vazyme) on a real-time PCR system. Each reaction was carried out in a total volume of 20 µL containing Taq Pro Universal SYBR qPCR Master Mix, 0.4 µM of each primer, and diluted DNA template. The primer pair D-spen-F (5′-GAAGTGGAACCGTACCAGGG-3′) / D-spen-R (5′-TTGCGCATTGCCTTTACCAC-3′) amplified an endogenous region of the *spen* locus that was unaffected by the editing constructs and served as a genomic reference to normalize differences in DNA input. The primer pair gs_F (5′-CTTGAAAAAGTGGCACCGAGTC-3′) / y-HA-R (5′-GATGGGTGGGAAATAGATGGGG-3′) was designed with one primer annealed to the inserted split-gRNA cassette and the other annealed to adjacent genomic sequence at the *yellow* locus. The resulting amplicon spanned the junction between the inserted cassette and the genome, enabling detection of HDR-mediated integration products. The thermal cycling conditions were as follows: initial denaturation at 95 °C for 30 s, followed by 40 cycles of 95 °C for 10 s and 65 °C for 60 s. Melting curve analysis was performed at the end of each run to verify amplification specificity. All reactions were performed with at least four technical replicates. The median Cp values of the technical replicates were used for subsequent calculations. For each biological replicate, ΔCp was calculated as the median Cp of the target amplicon minus that of the reference amplicon, and relative HDR efficiency was calculated using the 2^-ΔCp^ method. Statistical significance was assessed using ΔCp values by ordinary one-way ANOVA followed by Dunnett’s multiple-comparisons test, with standard Cas9 (with no fusion protein) serving as the control group.

### Generation of transgenic lines

To generate Cas9 lines, donor plasmids (500 ng/µL), gRNA helper plasmid BHDabg1^28^ (100 ng/µL), and the Cas9-expressing plasmid TTChsp70c9^29^ (450 ng/µL) were co-injected into *w^1118^*embryos by UniHuaii and Fungene Biotech. Injected individuals were crossed to *w^1118^* flies, and progeny carrying the drive element were identified by fluorescence. After confirming the insertion site by Sanger sequencing of genomic DNA, homozygous lines were established by inbreeding for several generations followed by crosses with *w^1118^* to confirm. The gRNA lines targeting *yellow*, *cinnabar*, *RpL35A*, *stall*, and *hairy* were previously generated^19,27,30,31^. Previously constructed Cas9 promoter lines include SNc92nG, SNc9sp2nG, SNc9sp3nG, and SNc9Cyp6nG^27^.

### Fly rearing and phenotypes

Flies were reared on modified Cornell standard cornmeal medium, with modifications including 10 g agar instead of 8 g per liter, the addition of 5 g soy flour, and omission of phosphoric acid. Flies were maintained at 25 °C under a 14/10 h day/night cycle and 60% humidity as described in previous reports^15,27^. To screen for fluorescence, flies were anesthetized using CO₂ and phenotyped using NIGHTSEA adapter SFA-GR for DsRed and SFA-RB-GO for EGFP. Fluorescent markers were expressed in the eyes of the white-eyed *w^1118^* background under the control of the 3×P3 promoter and the SV40 3′ UTR/terminator. EGFP and DsRed expression indicate the presence of the Cas9 and the gRNA drive allele, respectively.

In the *yellow* system, disruption of *yellow* (*y*) function causes a recessive yellow body phenotype resulting from reduced dark pigment in adults. Body color was therefore used to infer resistance genotypes. Because *y* is X-linked and recessive, males carrying a disrupted *y* allele are yellow due to hemizygosity, whereas females display the yellow phenotype only when both *y* copies are disrupted. Resistance alleles are classified as functional or nonfunctional, with only nonfunctional alleles expected to lose *y* activity. In addition, post-zygotic cleavage and end-joining can occur in embryos, producing additional resistance alleles in all or some cells of new individuals. In these situations, individuals were scored as "mosaic" if the yellow phenotype was present in only part of the body. In the *cinnabar* system, individuals lacking wild-type alleles typically produce bright orange eyes, although this phenotype could not be assessed in the *w^1118^* background because these flies lack eye pigmentation.

### Drive performance assays

Males from different Cas9 lines were crossed with females from various gRNA lines to generate male or virgin female progeny carrying both DsRed and EGFP fluorescent markers. Male Cas9 lines were used as grandparents in cross design to minimize potential maternal Cas9 deposition, which could provide additional Cas9 and affect the drive performance assay.

To account for variation among biological replicates, statistical analyses were performed in R using generalized linear mixed-effects models (GLMMs) implemented with the lme4 package^11,19^. Binary outcomes were analyzed using binomial GLMMs, where individual progeny were classified as either drive or non-drive (or other binary phenotypic categories depending on the analysis). For each experiment, vial ID was included as a random effect to account for replicate variation. Estimated marginal means and their standard errors were calculated using the emmeans package to obtain model-based expected proportions. The program used is available on Github (https://github.com/jchamper/Fusion-Cas9).

In the *yellow* system, each Cas9 fusion construct was compared with the corresponding standard Cas9 control using Wald z-tests with Dunnett adjustment. When zero counts or unstable model fitting occurred, two-sided Fisher’s exact tests with Holm correction were used. In the *RpL35A*, *stall*, *hairy*, and *cinnabar* systems, prespecified pairwise comparisons between Cas9 constructs were performed separately for crosses involving female and male drive parents, with *P* values calculated using Wald z-tests and adjusted for multiple comparisons using the Holm method.

## Results

### Initial Cas9 fusion protein screening reveals limited enhancement of HDR efficiency in Drosophila S2 cells

To investigate the influence of Cas9 fusions on HDR efficiency, we selected fusion proteins spanning multiple stages of the repair process, including end resection (CtIP and UL12) ^22,32,33^, recombination filament formation and stabilization (Spn-A and Brex27)^34^, DNA synthesis-associated repair (POLD3)^24^, regulation of chromatin accessibility (PRDM9)^26^, and cell-cycle restriction (Geminin degron)^22^. These repair-associated proteins were fused to either the N- or C-terminus of Cas9 via short flexible linkers, and nuclear localization signals (NLSs) were also included (756873v21A). The expression of fusion proteins was controlled by the constitutive promoter *Act5C* to maximize DNA cleavage in *Drosophila* S2 cells. A standard screening assay was applied to enrich positively transfected cells and systematically compare HDR efficiency from different Cas9 fusion proteins (Fig.1C). By introducing Cas9 cleavage at the wild-type *yellow* locus and supplying a donor template, HDR efficiencies mediated by distinct Cas9 fusion constructs could be compared by evaluating relative copy numbers of HDR-integrated drive alleles (Fig.1B).

**Fig. 1:**
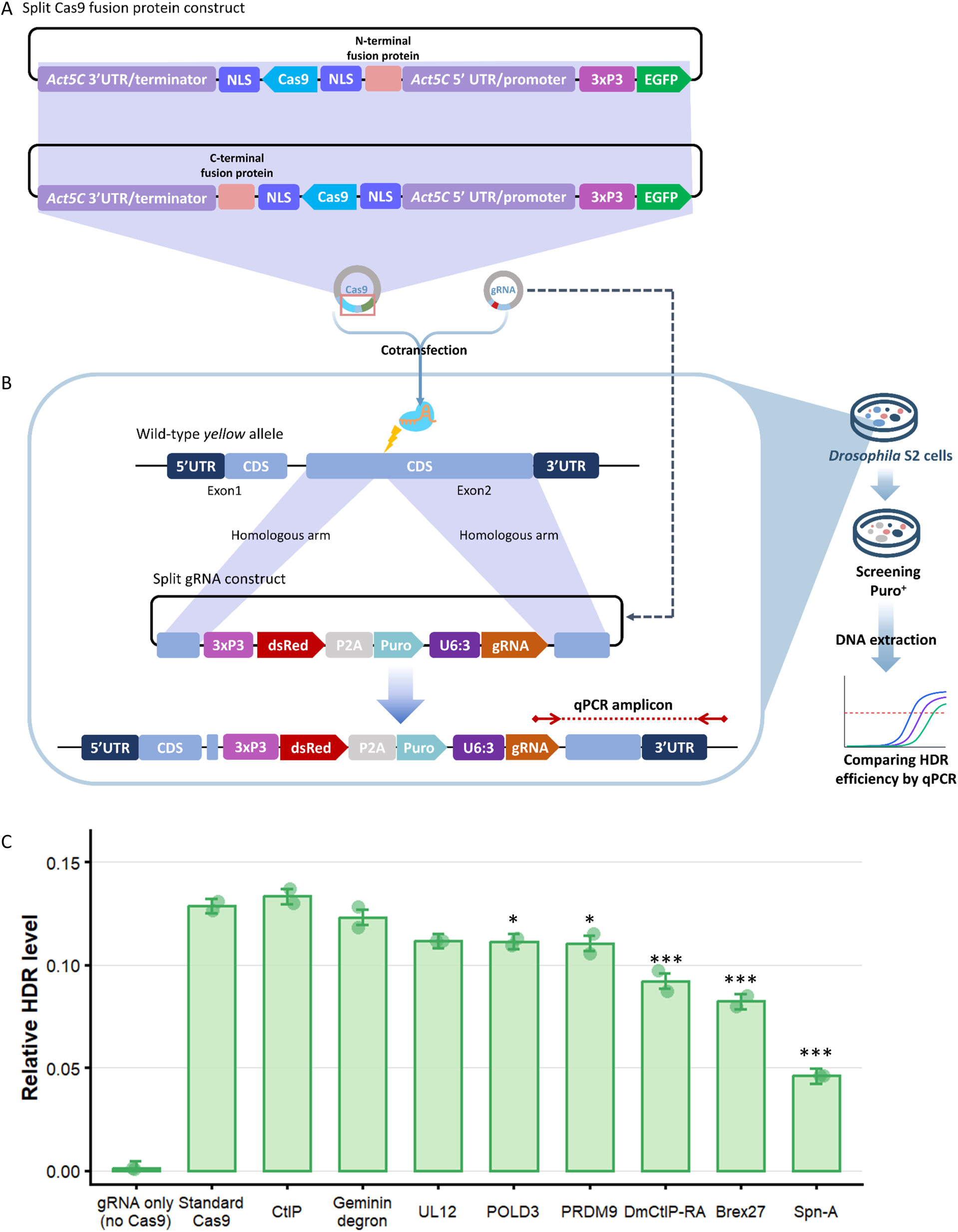
In *vitro* evaluation of HDR efficiency at the *yellow* locus with Cas9 fusions. **A.** Cas9 fusion protein expression constructs. Repair-associated proteins are fused to either the N- or C-terminus of Cas9 via short flexible peptide linkers or nuclear localization signals. Cas9 and its fusion proteins are expressed under the control of the upstream and downstream regulatory elements of *Act5C*. EGFP was included as a fluorescent reporter. **B.** gRNA construct, HDR-mediated integration, and experimental workflow for evaluating Cas9 fusion proteins in *Drosophila* S2 cells. Cas9 fusion and gRNA plasmids were cotransfected into S2 cells. The donor plasmid contains homology arms derived from the wild-type *yellow* locus, a *yellow*-targeting gRNA driven by the U6:3 promoter, a DsRed fluorescent marker, and a puromycin resistance cassette. Cas9/gRNA-mediated cleavage generates a cut at the *yellow* locus, enabling integration of the donor sequence through HDR. Following puromycin selection, genomic DNA was extracted from enriched cells and HDR efficiency was quantified by qPCR. **C.** HDR efficiency analysis of different Cas9 fusion proteins. “Standard” refers to Cas9 without fusion proteins. HDR levels were quantified by qPCR and normalized to a reference locus. Bars indicate the mean, and error bars denote standard deviation. Individual points represent independent replicate assays (n = 2). *P* values were calculated from ΔCt values using ordinary one-way ANOVA followed by Dunnett’s multiple comparisons test, with standard Cas9 as the control group. The gRNA-only group was included as a negative control. Statistically significant differences with standard Cas9 are marked with * (*P* < 0.05) and ***(*P* < 0.001). Source data are provided in Source Data S1.

We performed qPCR to assess relative HDR level (Fig.1C). The negative control transfected with gRNA plasmid alone produced negligible signal, indicating good assay specificity. Compared to standard (no fusion) Cas9, CtIP, Geminin degron, and UL12 fusions showed comparable HDR efficiencies, whereas POLD3 and PRDM9 fusions resulted in a modest but significant reduction. In contrast, DmCtIP-RA, Brex27, and Spn-A markedly reduced HDR efficiency, with Spn-A showing the strongest inhibitory effect. These results indicate that, in this S2 cell assay, most tested repair-associated fusions did not enhance HDR and often impair Cas9-mediated editing or repair outcomes.

### Cas9-POLD3 and Cas9-Geminin degron improve distinct aspects of drive performance across regulatory elements

Based on the cell experiments, we generated transgenic *Drosophila* lines carrying selected Cas9 fusion constructs that showed comparable performance in the *in vitro* assay and evaluated their performance in a split drive system targeting *yellow* (Fig.2A-B). The drive allele carried a gRNA targeting *yellow* and a fluorescent marker, enabling quantification of drive element inheritance and the formation of germline and embryo resistance alleles (Fig.2B, Fig.S1A and Fig.S2A). The dual-NLS Cas9 and corresponding fusion constructs used in the cell experiments were expressed in the germline using the *nanos* regulatory element to assess the effects of each fusion protein on drive performance *in vivo* (Fig.2A).

**Fig. 2:**
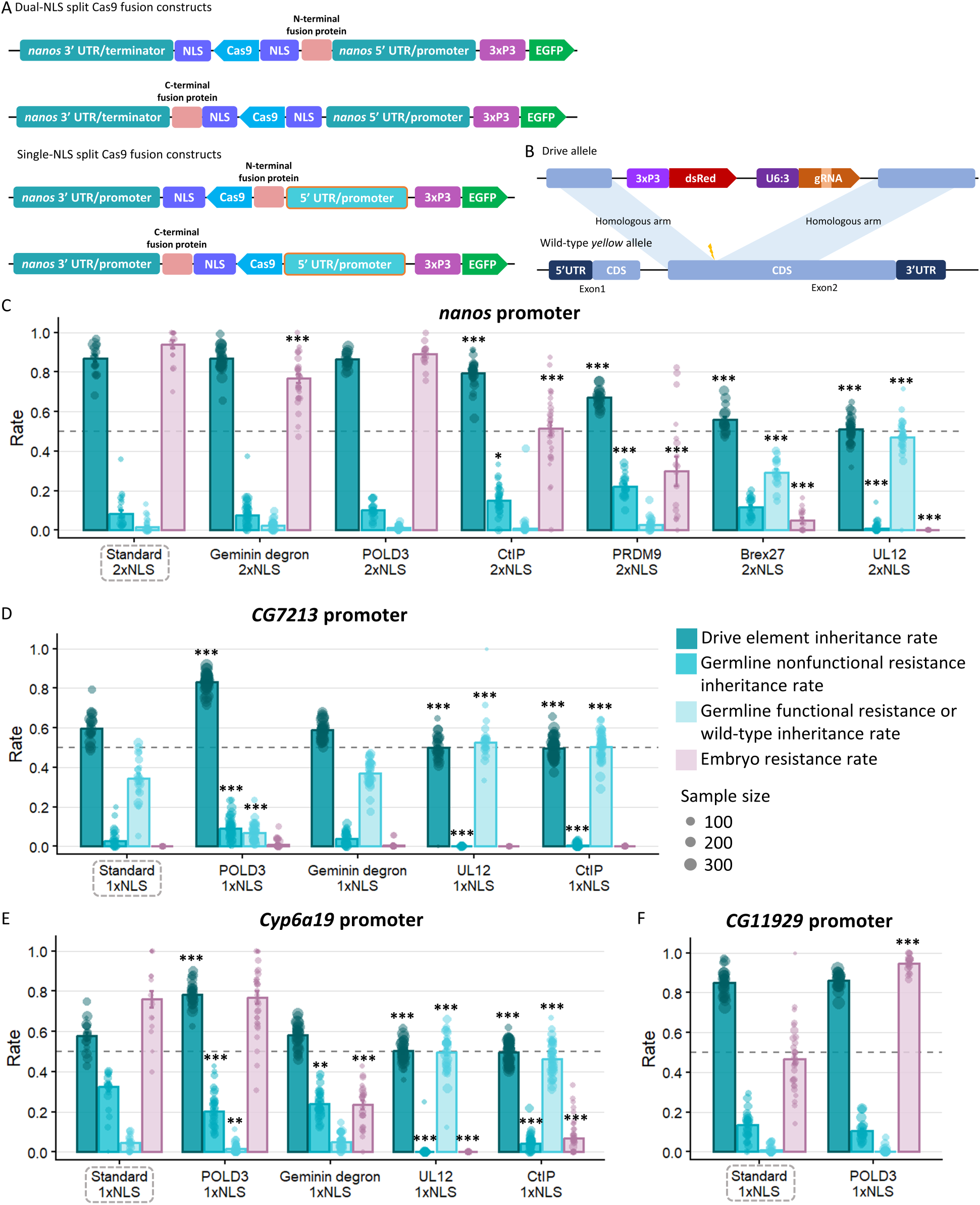
In *vivo* evaluation of Cas9 fusion proteins in a split drive system targeting *yellow*. **A.** Dual-NLS and Single-NLS split Cas9 fusion protein expression constructs. Cas9 and its fusion proteins are expressed under the control of the upstream and downstream regulatory elements of *nanos*. EGFP was included as a fluorescent reporter controlled by 3×P3 promoter and SV40 terminator. Purple boxes indicate NLS sequences (Nucleoplasmin or SV40). **B.** Schematic of the split gene drive targeting *yellow*. The drive allele contains a DsRed marker driven by 3×P3 and a gRNA driven by U6:3. The gRNA in the drive element directs Cas9 to cleave exon 2 of *yellow*, and the drive element is integrated into the *yellow* locus by HDR. Germline and embryo drive performance of dual-NLS Cas9 fusion constructs under the **C** *nanos* **D** *CG7213* **E** *Cyp6a19* **F** *CG11929* 5′ regulatory elements. Data corresponding to construct names enclosed in gray dashed boxes were obtained from a previous study^27^. Each data point represents an independent cross, and point size indicates the number of progenies scored. Error bars indicate standard error of the mean. Statistical significance relative to the corresponding standard Cas9 control was assessed using binomial generalized linear mixed models followed by Dunnett-adjusted comparisons. Fisher’s exact tests with Holm correction were used for comparisons involving zero counts or unstable model fits. \**P* < 0.05; \*\**P* < 0.01; \*\*\**P* < 0.001. Source data are provided in Source Data S2.

Under the *nanos* promoter, standard Cas9 exhibited high drive inheritance together with a high level of embryo resistance (Fig.2C, Fig.S3A). Cas9-POLD3 showed similar drive performance, maintaining both high drive inheritance and embryo resistance rate. In contrast, the CtIP, PRDM9, Brex27, and UL12 fusions reduced drive inheritance, germline nonfunctional resistance, and embryo resistance to varying degrees. This potentially demonstrates an overall reduction in effective Cas9 cleavage activity with these fusions. Notably, Cas9-Geminin degron maintained germline cleavage and drive inheritance at levels comparable to those of standard Cas9, while reducing embryo resistance rate.

Because the dual-NLS Cas9 driven by *nanos* already induced high drive inheritance, the scope for further improvement in this context was likely limited. Therefore, a lower activity Cas9 promoter was selected to further investigate the fusion proteins. Based on our previous study^27^, regulatory elements from three additional germline genes (*CG7213*, *Cyp6a19*, and *CG11929*) with distinct baseline drive performances were applied together with a single-NLS to further reduce Cas9 baseline activity without affecting the fusion protein itself (Fig.2A).

Under the *CG7213* regulatory element, standard Cas9 displayed a low level of drive inheritance, and embryo resistance was nearly zero (Fig.2D, Fig.S3B), consistent with relatively limited overall Cas9 activity. Compared with standard Cas9, Cas9-POLD3 significantly increased drive inheritance (*P <* 0.0001) and overall germline cleavage rates (*P <* 0.0001) while maintaining an almost undetectable level of embryo resistance. Cas9-Geminin degron showed drive inheritance and resistance allele formation broadly similar to those of standard Cas9. In contrast, drive inheritance for UL12-Cas9 and Cas9-CtIP remained close to the expected Mendelian frequency of 50%, with neither construct showing detectable cleavage (Fig.S3B).

Under the *Cyp6a19* regulatory element, standard Cas9 exhibited high germline cleavage activity, but repair outcomes were not predominantly directed toward drive conversion (Fig.S3C). Drive inheritance was moderate, while a substantial proportion of germline nonfunctional resistance alleles were generated (Fig.2E, Fig.S3C). Relative to standard Cas9, Cas9-POLD3 significantly increased drive inheritance (*P <* 0.0001) and reduced the formation of germline nonfunctional resistance alleles (*P <* 0.0001), while maintaining a similarly high level of embryo resistance (Fig.2H). In addition, Cas9-Geminin degron showed an overall germline drive performance similar to that of standard Cas9 but markedly reduced the embryo resistance rate. UL12-Cas9 and Cas9-CtIP again failed to produce detectable super-Mendelian inheritance. With the *CG11929* promoter, standard Cas9 and Cas9-POLD3 showed similarly high drive inheritance, but Cas9-POLD3 produced substantially higher embryo resistance allele formation (Fig.2F, Fig.S3D), again suggesting that this fusion produced higher cleavage activity in general.

Overall, the effects of Cas9 fusion proteins on drive performance were strongly dependent on the regulatory element. Cas9-POLD3 increased drive inheritance under the *CG7213* promoter, where baseline Cas9 activity was relatively low, and under the *Cyp6a19* promoter, where an increased proportion of germline cleavage events resulted in drive conversion. Notably, Cas9-POLD3 driven by *CG7213* combined high drive inheritance with nearly undetectable embryo resistance, identifying a promising construct for further evaluation in other drive systems. In contrast, Cas9-POLD3 was not superior under the *nanos* and *CG11929* promoters, in which drive inheritance was already high. Cas9-Geminin degron generally preserved germline drive performance across multiple regulatory elements while reducing the embryo resistance rate. In addition, the other fusion proteins reduced both germline and embryo drive activity, suggesting that these fusions may impair overall Cas9 activity.

### The effects of Cas9-POLD3 and Cas9-Geminin degron on drive performance depend on sex and drive system

Given that Cas9-POLD3 and Cas9-Geminin degron could improve drive performance in the previous drive system targeting *yellow*, we further evaluated these fusion constructs across different target genes and drive systems. Because the earlier experiments assessed only female drive performance in the *yellow* system, we further evaluated the effects of these fusion proteins across additional drive systems in both females and males (Fig.S2B).

In the drive targeting the eye pigment gene *cinnabar*^27^, we evaluated drive inheritance from female and male parents carrying either standard Cas9 or Cas9-POLD3 driven by the *CG7213* or *Cyp6a19* regulatory element. In both contexts, Cas9-POLD3 significantly increased female drive inheritance (*CG7213*, *P* = 0.000124; *Cyp6a19*, *P* < 0.0001) but had only a limited effect in males (Fig.3A-B), indicating that its enhancing effect in this system was potentially sex dependent.

**Fig. 3:**
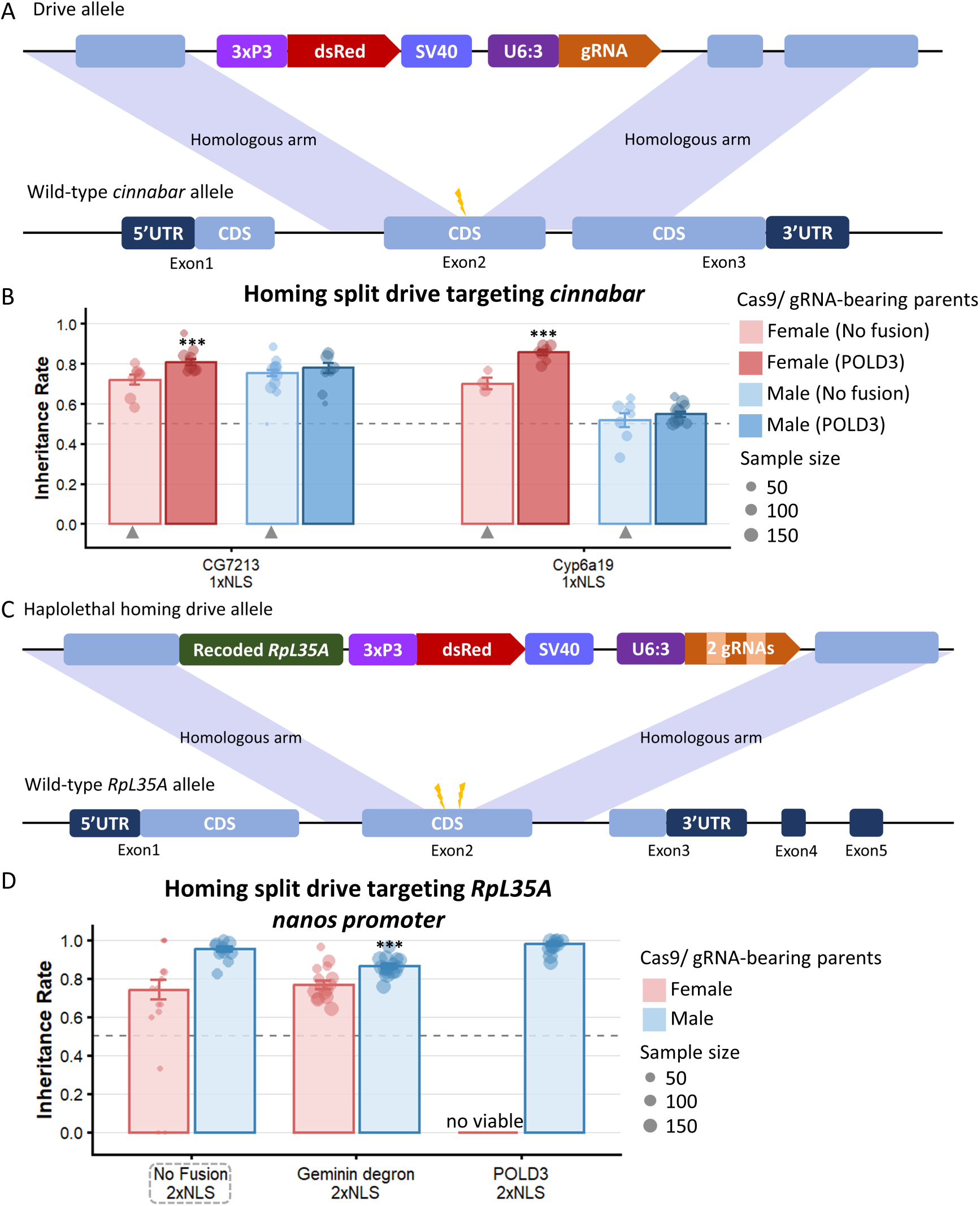
The evaluation of Cas9 fusion proteins in a split drive system targeting *cinnabar* and *RpL35A.* **A.** Schematic of the gRNA construct targeting *cinnabar*. The drive allele contains a DsRed marker driven by 3×P3 and a U6:3-driven gRNA targeting exon 2 of *cinnabar*. **B.** Drive inheritance in the *cinnabar* drive system. Data marked with gray triangles was obtained from our previous study^27^. Source data are provided in Source Data S3. **C.** Schematic of the gRNA construct targeting *RpL35A*. Two gRNAs are driven by U6:3 and target exon 2 of *RpL35A*. **D.** Drive inheritance in the *RpL35A* drive system. No viable progeny were obtained from female parents carrying Cas9-POLD3. Data corresponding to construct names enclosed in gray dashed boxes were obtained from a previous study^35^. Each data point represents an independent cross, and point size indicates the number of progenies scored. Error bars indicate standard error of the mean. Statistical significance relative to the corresponding standard Cas9 control was assessed using binomial generalized linear mixed models followed by Dunnett-adjusted comparisons. \*\*\**P* < 0.001. Source data are provided in Source Data S4.

In the rescue drive targeting the haplolethal gene *RpL35A*^19^, only one copy of the intact gene was insufficient for survival. Thus, embryo resistance is expected to render individuals nonviable. We found that female parents carrying Cas9-Geminin degron produced significantly more viable progeny than those carrying standard Cas9 (Fig.3D, *P* < 0.0001). This result may reflect reduced maternal Cas9 deposition, consistent with our earlier observation that Cas9-Geminin degron reduced embryo resistance formation. In contrast, female parents carrying Cas9-POLD3 produced no viable progeny, suggesting that this construct likely increased embryo Cas9 activity or maternal deposition compared to standard Cas9, thereby causing more embryonic lethality. This result was also consistent with the increased embryo resistance associated with Cas9-POLD3 driven by *CG11929* in the *yellow* system.

In the homing drive targeting the female fertility gene *stall*^31^, Cas9-POLD3 and Cas9-Geminin degron maintained high drive inheritance in both female and male parents but showed a slight reduction compared with standard Cas9 (Fig. S4). In the TARE drive targeting the essential but haplosufficient gene *hairy*^18^, Cas9-POLD3 showed inheritance rates comparable to standard Cas9, but Cas9-Geminin degron reduced drive inheritance among the progeny of females (Fig. S5).

Overall, these results indicate that Cas9-POLD3 and Cas9-Geminin degron exhibited distinct but generally consistent functional trends across different systems. Cas9-POLD3 increased drive inheritance, whereas its beneficial effect was limited in systems that already displayed high inheritance rates. In contrast, Cas9-Geminin degron consistently reduced embryo resistance formation, while its effects on drive inheritance varied depending on the drive architecture.

## Discussion

In this study, we combined a *Drosophila* S2 cell-based HDR assay with multiple *in vivo* drive systems to systematically evaluate the effects of DNA repair-related Cas9 fusion proteins on drive performance. Although none of the fusion proteins tested significantly enhanced HDR efficiency in S2 cells, *in vivo* analyses showed that Cas9-POLD3 increased drive inheritance under specific germline regulatory contexts, and Cas9-Geminin degron could significantly reduce embryo resistance allele formation. Most other fusions provided no improvement or reduced drive performance. Together, these results indicate that the effects of individual fusion proteins were jointly influenced by the regulatory elements, drive architecture, target site, and parental sex, indicating that Cas9 fusion strategies are highly context dependent.

Previous studies have shown that the protein fusions can influence folding and stability, nuclear localization, steric accessibility, and interactions with endogenous DNA repair factors^22,36–38^. We therefore prioritized fusion configurations that had previously been reported to retain activity, including fusion orientation, linker design, protein isoform, and boundaries of the functional domain or fragment. These configurations were derived from different experimental systems, and the candidate proteins used in this study also varied in species of origin. DmCtIP-RA and Spn-A used *Drosophila* sequences, UL12 used a viral sequence, and the remaining candidates used the human sequences originally tested. Heterologous protein modules may be affected by protein folding, post-translational regulation, or compatibility with endogenous *Drosophila* DNA repair factors^39^. Nevertheless, the two modules that produced the most pronounced improvements in drive performance, POLD3 and the Geminin degron, were both human-derived, while other *Drosophila*-derived candidates showed no obvious advantage. These findings indicate that heterologous origin does not necessarily restrict the function of fusion proteins in *Drosophila*. Instead, the relevant functional modules may be sufficiently conserved across species to remain recognizable and functional when interacting with other proteins, or perhaps derive their activity by direct DNA interactions. Future studies could directly compare *Drosophila* homologs of POLD3 and Geminin degron with the corresponding human-derived constructs to determine whether protein sequences from the target species could further improve drive performance.

Most candidate fusion proteins failed to improve drive performance, and the CtIP, PRDM9, Brex27, and UL12 fusion constructs reduced the effective cleavage events to varying degrees. These findings indicate that DNA repair protein configurations capable of enhancing HDR in other cell types or species do not necessarily improve drive performance in *Drosophila*. While we cannot rule out a potential increase in relative HDR rate with these fusions, the dominant effect was clearly a reduction in Cas9 cleavage rates. One possible explanation is that tethering of these modules to Cas9 may impair fusion protein expression, stability, or nuclear localization, and may also interfere with Cas9 cleavage through steric hindrance or mismatched timing between Cas9 activity and repair factor recruitment^36,40–42^. Importantly, the magnitude of these fusion-associated effects appeared to depend on the Cas9 baseline activity. Stronger Cas9 expression could partially compensate for fusion impairment, whereas high baseline activity could also limit the ability to detect additional improvements by fusion proteins. In the initial experiments, the strong dual-NLS Cas9 systems used in S2 cells and the *Drosophila* germ cells exhibited limited dynamic range for detecting further enhancement. Therefore, subsequent experiments employed weaker Cas9 regulatory elements generating different baseline levels of drive performance, providing a broader window for evaluating fusion effects. Differences between *Drosophila* germ cells and the original experimental systems of fusion proteins, including regulatory environments and repair pathway interactions, may further contribute to these effects. Because these parameters were not directly measured, this interpretation remains inferential.

The S2 cell-based assay provides a convenient platform for rapid screening of multiple fusion constructs, but its ability to predict *in vivo* drive performance is limited. In this study, the S2 assay and *in vivo* systems showed broadly consistent trends, with Cas9-POLD3 and Cas9-Geminin degron exhibiting relatively higher activity, while several other fusion constructs showed limited or reduced performance. However, the correspondence between the two systems was not consistent across all constructs. This discrepancy is likely related to differences between the S2 cell and the *Drosophila* germline, which differ not only in cellular state and DNA repair environment, but also in Cas9 regulatory context and developmental timing. Importantly, the qPCR assay quantified the relative abundance of HDR-mediated integration products normalized to a genomic reference, which was designed primarily to compare relative HDR-associated signals among constructs rather than to measure absolute repair outcomes at the target site. Therefore, HDR performance measured at the cellular level may not fully represent overall drive performance *in vivo*.

Among all fusion proteins tested, Cas9-POLD3 exhibited the most pronounced improvement in drive performance. However, Cas9-POLD3 did not enhance performance across all drive systems, with its benefits being most apparent for systems in which the baseline drive performance was not already saturated. In these contexts, Cas9-POLD3 may not only increase effective Cas9 activity but also possibly increase the proportion of cleavage events that proceed to drive conversion. This observation is consistent with previous studies showing that POLD3 can promote both HDR and end-joining pathways^24^, but our study did not directly confirm an increase in the HDR rate.

Although POLD3 is considered a component of DNA polymerase δ and is involved in DNA synthesis during repair processes^43^, the POLD3 fragment used in the Cas9-POLD3 fusion in this study lacks the N-terminal region required for interactions with other DNA polymerase δ subunits^24^. Therefore, Cas9-POLD3 may enhance repair by modifying the local repair environment surrounding DNA double-strand breaks or by facilitating Cas9 release from the cleavage site, thereby promoting the initiation of subsequent repair processes. Alternatively, POLD3 fusion may enhance HDR indirectly by increasing effective Cas9 activity, which may increase the proportion of early germline cleavage events that occur during a more favorable repair window for HDR before meiosis II when HDR is no longer possible (Fig.S6). If cleavage of the wild-type allele is complete before meiosis II, then a further increase in activity (with the same expression pattern) would increase the fraction of alleles cut before the optimal repair window and thus actually reduce overall drive conversion efficiency. At the same time, high Cas9 activity may also increase maternal Cas9 deposition, resulting in increased embryo cleavage and the formation of embryo resistance alleles. Thus, POLD3 fusion may produce both beneficial effects on drive conversion and potential costs associated with increased embryo resistance, with the overall outcome determined by the specific Cas9 regulatory context. We note that Cas9 dissociation, cleavage kinetics, and the recruitment of DNA repair factors were not directly examined in this study, and therefore, this proposed mechanism requires further validation.

In the split drive system targeting *cinnabar*, Cas9-POLD3 produced a greater enhancement of drive inheritance in the female germline, whereas only a non-significant increase was observed in the male germline (Fig.3B). Female and male germ cells in *Drosophila* differ in their differentiation programs, cell cycle progression, and chromosome dynamics. In particular, male meiosis in *Drosophila* lacks canonical synaptonemal complex formation and meiotic crossing over, whereas the female germline undergoes conventional meiotic recombination^44,45^. Drive conversion can still occur in the male germline because Cas9 cleavage and homing may take place in premeiotic germ cells^14,46,27^. However, this suggests that developmental stages and DNA repair windows permissive for drive conversion may differ between sexes. Our previous work has shown that the same Cas9 construct can generate different degrees of inheritance bias in the female and male germlines^27^. In the *cinnabar* system, POLD3 may further enhance this biased drive conversion in the female germline, where cleavage timing may better coincide with HDR-preferred conditions.

While an increase in drive conversion rate would benefit gene drive in most organisms, many species are critically limited in overall Cas9 activity. These include *Culex* mosquitoes^47,48^, the diamondback moth^17^, mice^14,49,50^, and possibly *Aedes* mosquitoes^13,51,52^. Use of Cas9-POLD3 fusions could potentially substantially benefit gene drives trials in these organisms.

Unlike Cas9-POLD3, the principal advantage of Cas9-Geminin degron was its significant reduction of embryo resistance formation. It has been reported that the Geminin degron can restrict the temporal window of Cas9 activity through cell-cycle-dependent protein degradation^53,54^. This regulation may shorten the persistence of maternally deposited Cas9 after fertilization, thereby reducing the formation of embryo resistance alleles while largely preserving germline drive conversion. However, an alternative explanation is that, similar to other fusion proteins that can impair drive performance, the Geminin degron may simply reduce overall Cas9 activity. This is somewhat inconsistent with our results showing that even when germline activity was low, it was still preserved with this construct. However, the mechanism remains uncertain. Because Cas9 stability and maternal deposition were not directly measured in this study, these mechanistic hypotheses require further investigation.

In summary, this study shows that DNA repair-related Cas9 fusions can improve several aspects of drive performance, but their effects are conditional rather than universal. They depend not only on the functional properties of the fusion module but also on multiple context-dependent factors, including Cas9 expression patterns determined by the regulatory elements, sex, target site characteristics, and gene drive types. Cas9-POLD3 was most beneficial in contexts with low baseline drive performance or high levels of germline nonfunctional resistance formation, whereas Cas9-Geminin degron primarily reduced resistance allele formation from maternal deposition. The selection and application of fusion modules should therefore be guided by the primary limitation of the specific drive system. Future studies should evaluate Cas9-POLD3 and Cas9-Geminin degron across additional drive systems, target sites, regulatory contexts, and species to further determine their robustness. Systematic comparisons with different fusion orientations, linker designs, and Cas9 construct backgrounds, combined with direct measurements of protein expression, nuclear localization, cleavage activity, and repair outcomes, should help identify fusion strategies that enhance drive conversion or reduce resistance formation while clarifying their mechanisms.

## Acknowledgements

This study was supported by the National Natural Science Foundation of China (W2432018, 32270672, and 32302455), and the Center for Life Sciences.

## Supplemental Information

**Table S1.** List of donor plasmid names with Cas9 fusion protein.

| Plasmids used for cell-based HDR assays |  |  |  |  |
| --- | --- | --- | --- | --- |
| Plasmid name | 5' regulatory element (Length) | 3' regulatory element (Length) | Fusion protein (Fusion site) | NLS number |
| SNc92Act5G | *Act5C* (2482/1837) | *Act5C* (1571/121) | No Fusion | 2 |
| SNc92BrAct5G | *Act5C* (2482/1837) | *Act5C* (1571/121) | Brex27 (C terminal) | 2 |
| SNc92CtAct5G | *Act5C* (2482/1837) | *Act5C* (1571/121) | CtIP (C terminal) | 2 |
| SNc92DmAct5G | *Act5C* (2482/1837) | *Act5C* (1571/121) | DmCtIP-RA (C terminal) | 2 |
| SNc92geAct5G | *Act5C* (2482/1837) | *Act5C* (1571/121) | Geminin (C terminal) | 2 |
| SNc92POAct5G | *Act5C* (2482/1837) | *Act5C* (1571/121) | Pold3 (C terminal) | 2 |
| SNc92PRAct5G | *Act5C* (2482/1837) | *Act5C* (1571/121) | PRDM9 (N terminal) | 2 |
| SNc92SpAct5G | *Act5C* (2482/1837) | *Act5C* (1571/121) | Spn-A (C terminal) | 2 |
| SNc92ULAct5G | *Act5C* (2482/1837) | *Act5C* (1571/121) | UL12 (N terminal) | 2 |

| Plasmids used for *Drosophila* drive performance assays |  |  |  |  |
| --- | --- | --- | --- | --- |
| Plasmid name | 5' regulatory element (Length) | 3' regulatory element (Length) | Fusion protein (Fusion site) | NLS number |
| SNc9Cyp6POnG | *Cyp6a19* (1825/14) | *nanos* (839/78) | Pold3 (C terminal) | 1 |
| SNc9Cyp6CtnG | *Cyp6a19* (1825/14) | *nanos* (839/78) | CtIP (C terminal) | 1 |
| SNc9Cyp6genG | *Cyp6a19* (1825/14) | *nanos* (839/78) | Geminin (C terminal) | 1 |
| SNc9Cyp6ULnG | *Cyp6a19* (1825/14) | *nanos* (839/78) | UL12(N terminal) | 1 |
| SNc9sp2POnG | *CG7213* (1338/124) | *nanos* (839/78) | Pold3 (C terminal) | 1 |
| SNc9sp2CtnG | *CG7213* (1338/124) | *nanos* (839/78) | CtIP (C terminal) | 1 |
| SNc9sp2genG | *CG7213* (1338/124) | *nanos* (839/78) | Geminin (C terminal) | 1 |
| SNc9sp2ULnG | *CG7213* (1338/124) | *nanos* (839/78) | UL12 (N terminal) | 1 |
| SNc9sp3POnG | *CG11929* (1570/147) | *nanos* (839/78) | Pold3 (C terminal) | 1 |
| SNc92BrnG | *nanos* (672/261) | *nanos* (839/78) | Brex27 (C terminal) | 2 |
| SNc92CtnG | *nanos* (672/261) | *nanos* (839/78) | CtIP (C terminal) | 2 |
| SNc92genG | *nanos* (672/261) | *nanos* (839/78) | Geminin (C terminal) | 2 |
| SNc92POnG | *nanos* (672/261) | *nanos* (839/78) | Pold3 (C terminal) | 2 |
| SNc92PRnG | *nanos* (672/261) | *nanos* (839/78) | PRDM9 (N terminal) | 2 |
| SNc92ULnG | *nanos* (672/261) | *nanos* (839/78) | UL12 (N terminal) | 2 |

**Supplementary Figure 1.**
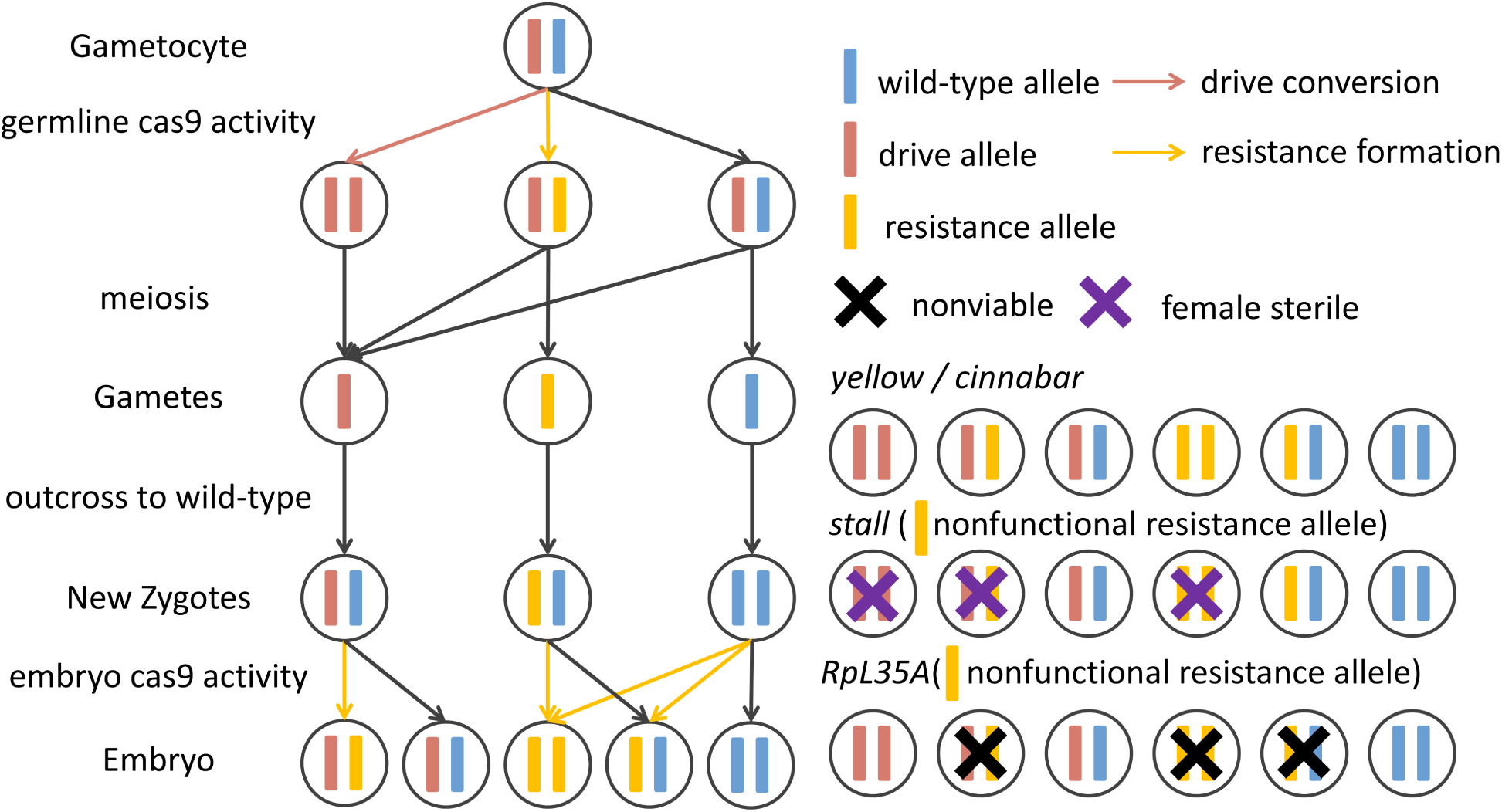
Schematic of inheritance and resistance formation of homing drive systems. In the germline of drive/wild-type heterozygotes, Cas9-mediated cleavage of wild-type alleles can be repaired by homology-directed repair, resulting in drive conversion, or by end-joining, generating a resistance allele. Uncut wild-type alleles may also be retained. Following meiosis and outcrossing to wild-type individuals, drive, resistance, and wild-type alleles are transmitted to the resulting zygotes. Maternal Cas9 deposition can further cleave wild-type alleles and generate resistance alleles in embryos. The genotype-specific viability and fertility outcomes for drives targeting *yellow*, *cinnabar*, *stall*, and *Rpl35A* are shown on the right, specifically for nonfunctional resistance alleles. Consistent with the properties of the target genes, all genotypes are viable in the *yellow* and *cinnabar* systems. Loss of both functional *stall* alleles causes female sterility, whereas fewer than two functional copies of *Rpl35A* (either drive, functional resistance, or wild-type) result in nonviability.

**Supplementary Figure 2.**
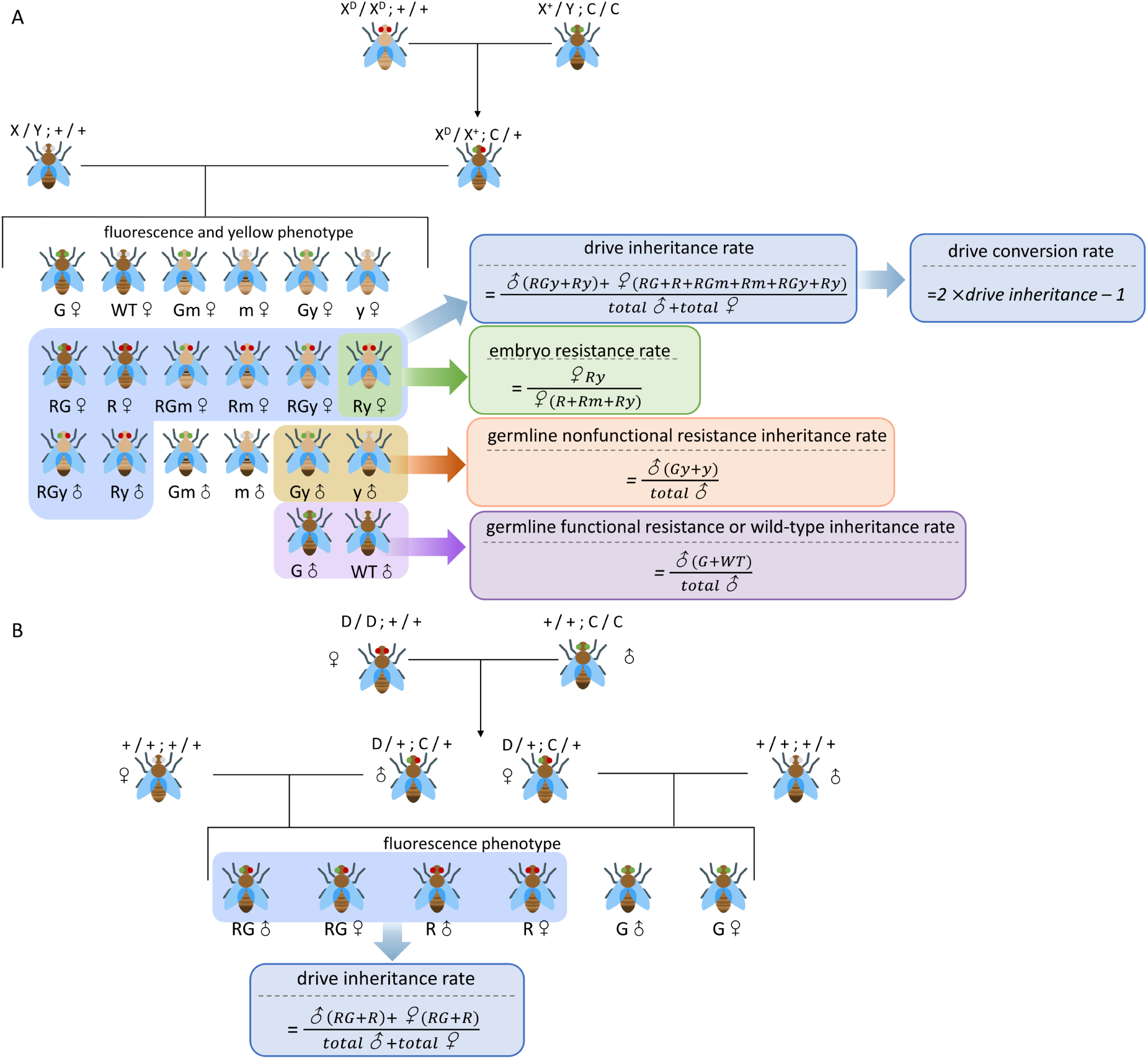
Cross scheme, phenotype classification, and calculation of gene drive performance parameters. **A.** Crossing scheme used to quantify drive performance in the split drive system targeting *yellow*. Progeny were classified according to sex, fluorescent marker, and body color phenotype, including DsRed (R), EGFP (G), yellow (y), mosaic for yellow (m), and wild type (WT). X^D^ refers to the drive allele, and C refers to the Cas9 allele. The drive inheritance rate and conversion rate were calculated based on the proportion of progeny carrying R. Embryo resistance was estimated from the proportion of Ry females among all females carrying R, reflecting resistance allele formation during embryo development. Germline nonfunctional resistance was estimated from the proportion of yellow-bodied male classes (including Gy and y males) among all males, reflecting nonfunctional resistance allele formation in the germline. Somatic expression was evaluated by the body color of female progeny from the initial cross carrying both fluorescent markers. More yellow pigmentation indicated higher somatic expression. Note that in the absence of somatic expression, embryo resistance can be calculated to include EGFP Cas9 females together with the phenotypes shown. Related data are provided in Source Data S2. **B.** Crossing scheme used to quantify drive performance in the split drive system targeting *cinnabar*, *RpL35A*, *stall*, and *hairy*. The drive inheritance rate was calculated based on the proportion of progeny carrying R.

**Supplementary Figure 3.**
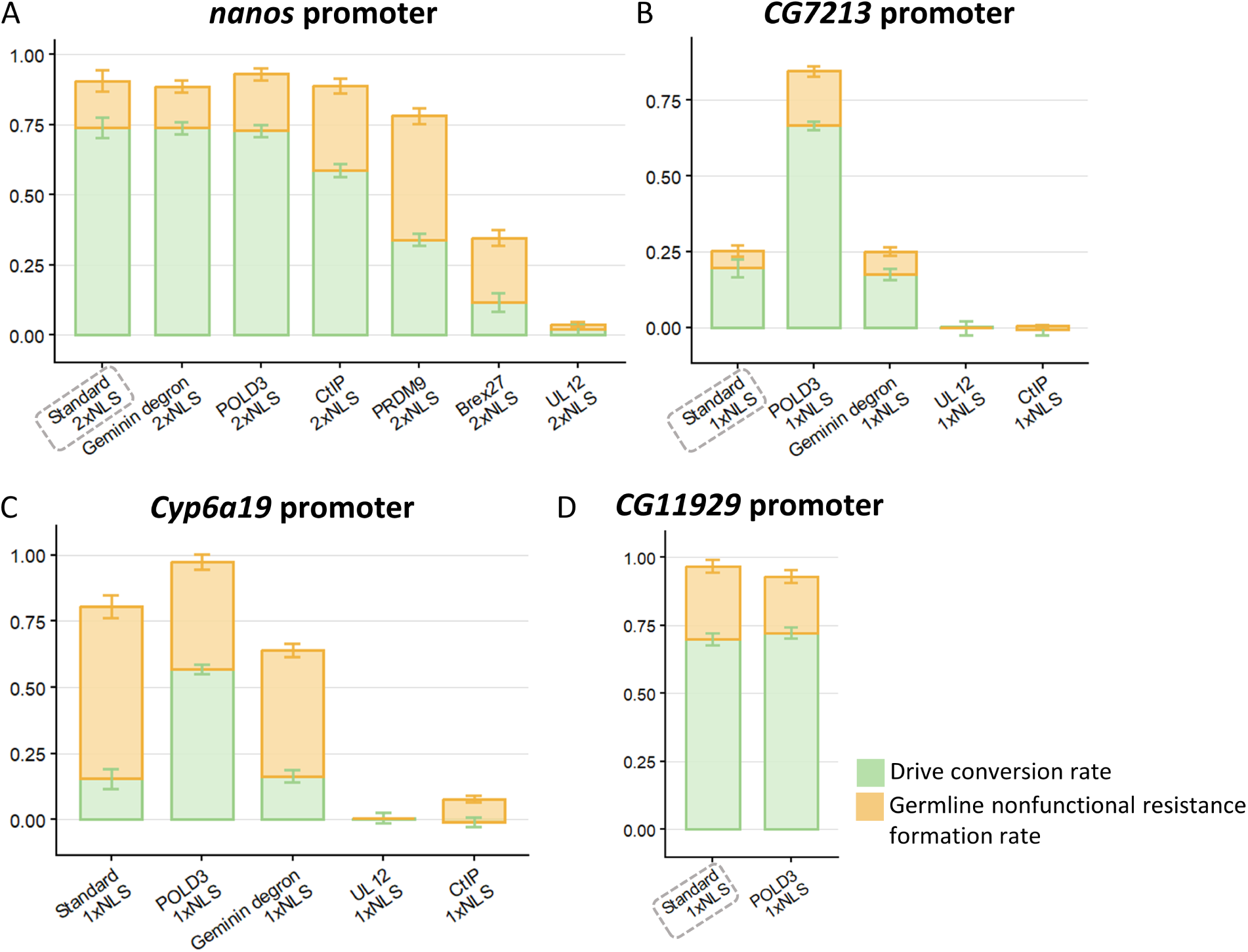
Drive conversion and germline nonfunctional resistance generated by Cas9 fusion constructs. Drive performance of fusion protein Cas9 constructs under the **A** *nanos* **B** *CG7213* **C** *Cyp6a19*, and **D** *CG11929* regulatory elements for Cas9. “Standard” refers to Cas9 without fusion proteins. NLS - Nuclear Localization Signal. Green and orange columns indicate the drive conversion rate and germline nonfunctional resistance rate, respectively. For improved visualization, drive conversion rates calculated as slightly below zero are displayed as zero. Data corresponding to construct names enclosed in gray dashed boxes were obtained from a previous study^27^. Error bars indicate standard error of the mean. Source data are provided in Source Data S2.

**Supplementary Figure 4.**
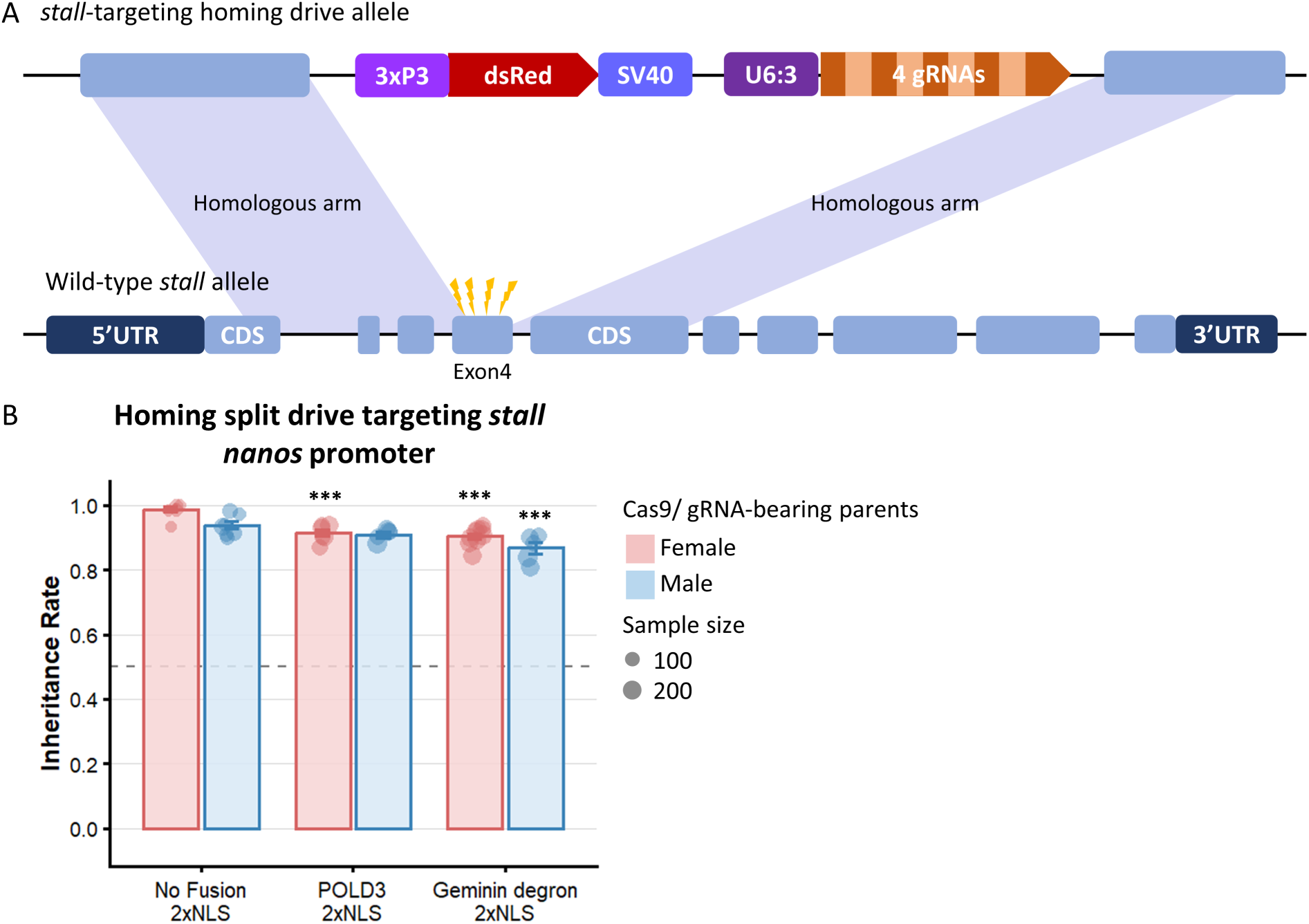
Drive performance in the homing split drive targeting *stall*. **A.** Schematic of the homing drive allele targeting *stall* and the wild-type *stall* allele. The drive allele contains a DsRed marker driven by 3×P3 and four gRNAs driven by U6:3. The gRNAs in the drive element direct Cas9 to cleave exon 4 of *stall*, and the drive element is then integrated into the *stall* locus by HDR. **B.** Drive inheritance from female and male parents carrying the drive allele targeting *stall* together with standard dual-NLS, Cas9-POLD3, or Cas9-Geminin degron driven by *nanos*. Each point represents an independent cross, and point size indicates the number of progeny scored. Error bars indicate standard error of the mean. \*\*\**P* < 0.001. Source data are provided in Source Data S5.

**Supplementary Figure 5.**
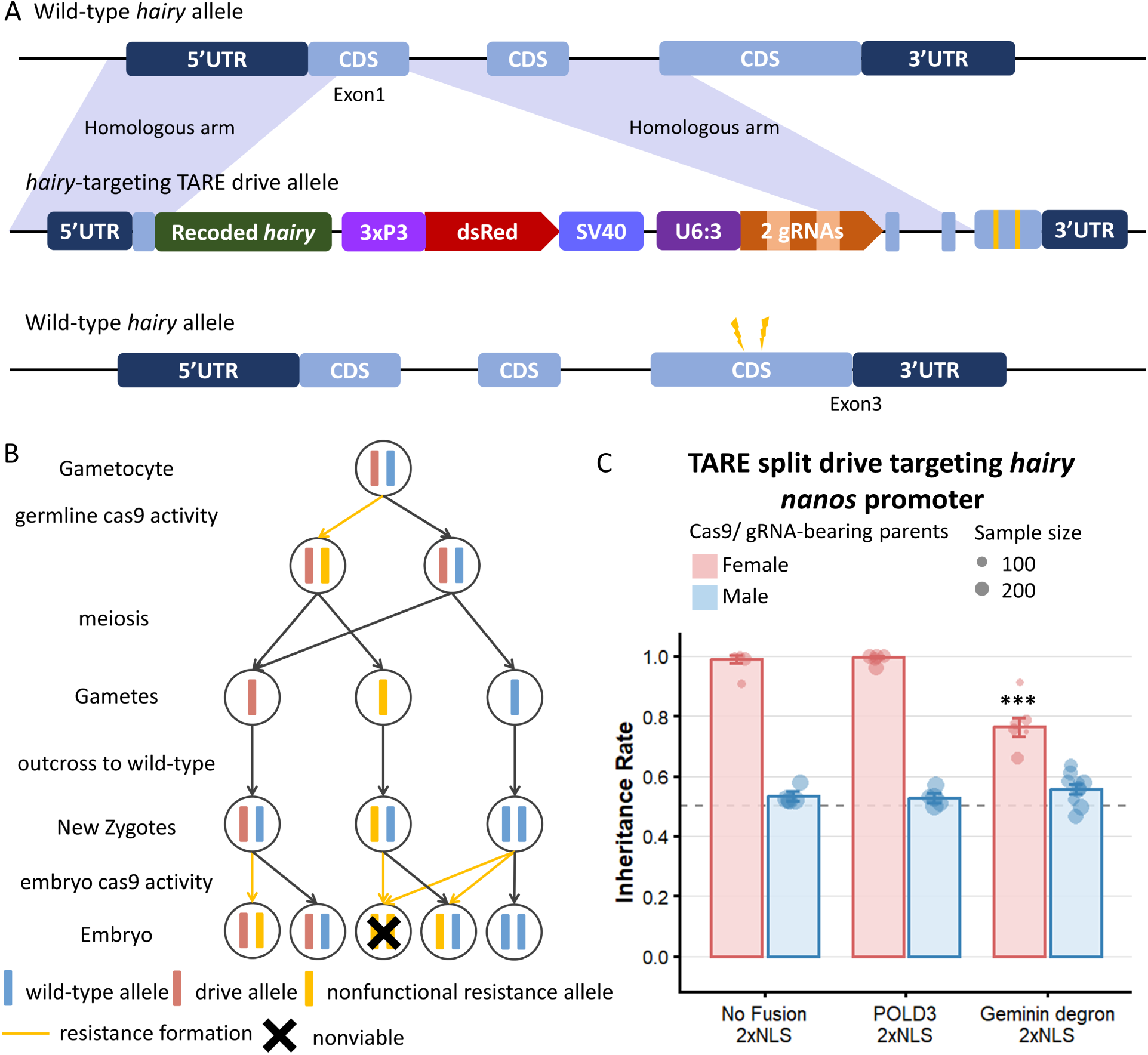
Drive performance in TARE split drive targeting *hairy*. **A.** Schematic of the TARE drive allele targeting *hairy* and the wild-type *hairy* allele. The TARE drive cassette is integrated into the exon 1 of *hairy* and contains a DsRed marker driven by 3×P3 and two gRNAs driven by U6:3. The gRNAs in the drive element direct Cas9 to cleave exon 3 of *hairy* and form disrupted/nonfunctional resistance alleles. **B.** Schematic of allele formation and transmission in a TARE drive. Germline Cas9 cleavage converts wild-type target alleles into disrupted alleles. Following meiosis and outcrossing, additional cleavage caused by embryo Cas9 activity can disrupt the remaining wild-type alleles. Progeny carrying two disrupted target alleles are nonviable, thereby increasing the relative inheritance of the drive among surviving progeny. **C.** Drive inheritance from female and male parents carrying the drive allele targeting *hairy* together with dual-NLS standard Cas9 or Cas9-POLD3 driven by *nanos*. Each point represents an independent cross, and point size indicates the number of progeny scored. Error bars indicate standard error of the mean. \*\*\**P* < 0.001. Source data are provided in Source Data S6.

**Supplementary Figure 6.**
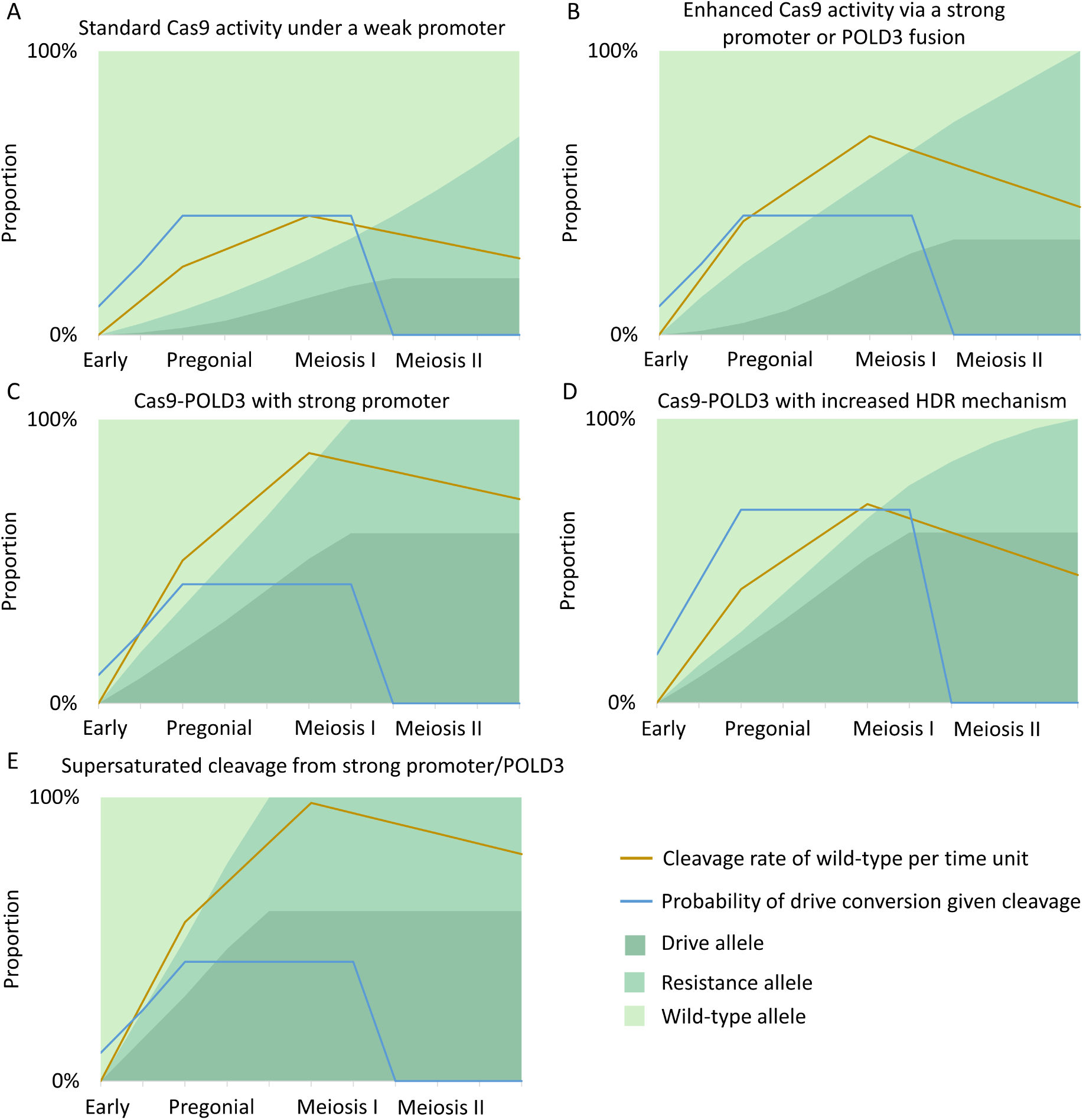
Hypothetical mechanisms by which Cas9-POLD3 fusion may enhance gene-drive performance. Charts show the cleavage rate and chance of successful drive conversion (vs. end-joining repair) if cleavage occurs. The background color shows the proportion of each (initially wild-type) allele. Proportions are intended to illustrate proposed mechanisms rather than be quantitatively precise. **A.** Standard Cas9 driven by a weak promoter, resulting in low cleavage activity and limited drive conversion. **B.** Enhanced Cas9 activity achieved through either a stronger promoter or fusion with POLD3, leading to increased cleavage and drive conversion. **C.** A scenario in which Cas9-POLD3 driven by a strong promoter further enhances drive performance primarily through increased cleavage activity. Because cleavage of wild-type alleles is completed before meiosis II (when HDR is not possible), a higher proportion of cut alleles are converted to drive alleles. **D.** An alternative scenario in which Cas9-POLD3 enhances drive performance by increasing drive conversion following cleavage, rather than further enhancing the cleavage rate. **E.** A scenario in which cleavage activity approaches saturation early in germline development (cleavage rates are higher than part C) under a strong promoter or POLD3 fusion, reducing drive conversion because more cleavage occurs before the optimal window.

